# Orphan cytochrome P450 20A1 CRISPR/Cas 9 mutants and neurobehavioral phenotypes in zebrafish

**DOI:** 10.1101/2021.07.23.453406

**Authors:** Nadja R. Brun, Matthew C. Salanga, Francisco X. Mora-Zamorano, David C. Lamb, Jared V. Goldstone, John J. Stegeman

**Affiliations:** Biology Department, Woods Hole Oceanographic Institution, Woods Hole, MA, 02543, USA; Department of Biological Sciences, Northern Arizona University, Flagstaff, AZ, 86011, USA; Faculty of Health and Life Sciences, Swansea University, Swansea, SA2 8PP, UK

**Author notes:** These authors contributed equally to this work (Nadja R. Brun, Matthew C. Salanga).

**Keywords:** vertebrate, cytochrome P450, anxiety, hyperactivity, mental health disorder, *Danio rerio*

## Abstract

Orphan cytochrome P450 (CYP) enzymes are those for which biological substrates and function(s) are unknown. Cytochrome P450 20A1 (CYP20A1) is the last human orphan P450 enzyme, and orthologs occur as single genes in every vertebrate genome sequenced to date. The occurrence of high levels of *CYP20A1* transcripts in human substantia nigra and hippocampus and abundant maternal transcripts in zebrafish eggs strongly suggest roles both in the brain and during early embryonic development. Patients with chromosome 2 microdeletions including *CYP20A1* show hyperactivity and bouts of anxiety, among other conditions. Here, we created zebrafish *CYP20A1* mutants using CRISPR/Cas9, providing vertebrate models with which to study the role of *CYP20A1* in behavior and other neurodevelopmental functions. The homozygous *cyp20a1* null mutants exhibited significant behavioral differences from wild-type zebrafish, both in larval and adult animals. Larval *cyp20a1*−/− mutants exhibited a strong increase in light-simulated movement (i.e., light-dark assay), which was interpreted as hyperactivity. Further, the larvae exhibited mild hypoactivity during the adaptation period of the optomotor assays. Adult *cyp20a1* null fish showed a pronounced delay in adapting to new environments, which is consistent with an anxiety paradigm. Taken together with our earlier morpholino *cyp20a1* knockdown results, the results described herein suggest that the orphan CYP20A1 has a neurophysiological role.

## INTRODUCTION

Cytochromes P450 (CYP; P450), a superfamily of enzymes found in every branch of life, catalyze a vast array of oxidation reactions, as well as the reduction and rearrangement of endogenous and exogenous compounds [1]. In vertebrates, including humans, CYP enzymes catalyze both physiological and toxicological reactions and play critical roles in many developmental stages.

When the physiological substrate(s) and function of a CYP are unknown, it is defined as an “orphan” P450. The functions of the majority of human and (by extrapolation) other mammalian P450s are known, although a few remain mysterious despite decades of intensive research [2–4]. Notable among these orphan CYPs is CYP20A1, the sole member of the CYP20 family, found in a single copy in all vertebrate genomes sequenced to date. CYP20A1 is the last human orphan P450 for which no biological or catalytic function is known.

While the activity of recombinant human CYP20A1 has been tested with possible substrates, no oxidation reaction was found to occur with steroids or selected biogenic amines [5]. Likewise, the activity of recombinant zebrafish Cyp20a1 has been tested with several different substrates without success [6]. Recently, human CYP20A1 expressed in yeast was observed to be weakly active with luminogenic substrates, as well as aniline [7, 8], suggesting that endogenous substrates may yet be identified.

Tissue and organ-specific expression patterns of enzymes such as *CYP20A1* can provide insights into function. In humans, the expression of *CYP20A1* transcripts varies in an organ-dependent manner. Expression is especially abundant in the hippocampus and substantia nigra regions of the brain [5], regions that are predominantly associated with learning and memory, and which are involved in neurodegenerative diseases including hyperactivity disorders (e.g., ADHD), panic disorders, social anxiety, and bipolar disorders. Such disorders affect >10% of the global population (~748 million people) [9, 10]. In other vertebrates, high levels of *cyp20a1* transcript occur in the brain and gonads of adult zebrafish [6] as well as in unfertilized eggs [11] and the notochord [12] of developing zebrafish, and during embryonic development of mice [13]. These findings suggest the participation of *CYP20A1* in vertebrate development, as well as its potential involvement in endocrine and neuronal processes.

We have previously demonstrated that transient morpholino knockdown of *cyp20a1* in zebrafish resulted in behavioral abnormalities, including increased latency or reduced responsiveness to a visual stimulus in larvae at 6 days post-fertilization (dpf). Morphants also exhibited a higher level of total physical activity and more bursts of movement than the control larvae; zebrafish behaviors that are consistently interpreted as hyperactivity [6]. Now we have developed *cyp20a1* mutant zebrafish to further interrogate the relationship between Cyp20a1 and behavioral phenotypes. Using CRISPR/Cas9, we generated zebrafish with lesions in the *cyp20a1* coding locus, resulting in a *cyp20a1*(−/−) crispant line following additional standard breeding. The *cyp20a1* crispants were examined for behavioral phenotypes in both larval and adult zebrafish. Ultimately, this *cyp20a1*(−/−) zebrafish may enable further characterization of genetic involvement in behavioral disorders, and the discovery of potential therapies at the molecular level.

## MATERIALS AND METHODS

### Animal husbandry

Experimental and husbandry procedures using zebrafish were approved by the Woods Hole Oceanographic Institution’s Animal Care and Use Committee. AB strain wild-type zebrafish were used in these studies. Embryos were obtained through pairwise or group breeding of adults using standard methods, rinsed with system water, and moved to clean polystyrene Petri dishes with 0.3X Danieau’s solution (17.4 mM NaCl, 0.21 mM KCl, 0.12 mM MgSO_4_, 0.18 mM Ca(NO_3_)_2_, and 1.5 mM HEPES at pH 7.6). Embryos were cultured at 28.5 °C and a 14 hr light – 10 hr dark diurnal cycle. The 0.3X Danieau’s solution was replaced at 24 hours post-fertilization (hpf) and all dead or defective embryos were removed. Larvae were fed daily with a diet according to their age starting with rotifers (*Brachionus rotundiformis*) at 5 days post-fertilization (dpf), then rotifers coupled with brine shrimp (*Artemia franciscana*) at 9 dpf, adding pellet feed (Gemma Micro 300, Skretting) at 21 dpf. The fish were then exclusively fed with brine shrimp and pellets from 30 dpf onward. To anesthetize the adult fish to obtain fin biopsies, the fish were immersed in fresh buffered Tricaine (3-amino benzoic acid ethyl ester; Sigma A-5040) diluted in system water (0.016%^w/v^) until motionless. Following fin biopsy, the adults were returned to their aquatic habitat and fed brine. The biopsied fish were allowed 7-10 days to recover before any additional handling.

### sgRNA site selection and synthesis

The coding sequence of exon 2 (reference sequence ZDB-GENE-030903-3) was queried for putative targets using the “CHOPCHOP” web tool [14]. Based on this analysis, we selected three targets opting for sequences that contained a G nucleotide within the first three nucleotides of the target sequence and no predicted off-target site.

Briefly, transcription was conducted using the MEGAscript (Ambion, AM1330) or MAXIscript (Ambion, AM1309) *in vitro* transcription reaction kits according to the manufacturer’s instructions using 80-200 ng of purified PCR products (see *PCR - sgRNA template preparation*). The samples were then incubated at 37°C between 4 and 5 hours; 80 ng of template DNA was used for the MAXIscript reaction and 200 ng of template DNA was used for the MEGAscript reactions.

### Microinjection equipment

Embryos were injected using a pneumatic microinjector (Model PV-820, World Precision Instruments). Injection needles were pulled from borosilicate capillary tubes (TW100F-4, WPI) using a vertical pipette puller (Model P-30, Sutter Instruments Inc.).

### Microinjection solutions

1-2 nl of injection solution was targeted to the yolk compartment of one-cell embryos immediately below the developing zygote. Injection solutions consisted of combinations of Cas9 recombinant protein (PNA Bio, CP-01) 1 μg μl^−1^, Cas9 mRNA (from Addgene plasmid #51307 [15]) 200-400 ng μl^−1^, H2B-RFP mRNA 200-400 ng μl^−1^, and pooled sgRNA 50-200 ng μl^−1^ (**Supplementary Table 1**).

### mRNA synthesis

1-5 μg of CS2-plasmid containing the ORF for Cas9 or H2B-RFP was linearized via Not1 endonuclease digestion followed by phenol:CHCl_3_:IAA extraction and EtOH precipitation. Next, 1 μg linearized plasmid was used as a template in the SP6 mMessage mMachine *in vitro* transcription reaction (Ambion, AM1344) according to the manufacturer’s instructions.

### PCR

Endpoint PCR for genotyping or single guide RNA template preparation was carried out using Q5 (M0491 NEB) or Taq (M0267 NEB) polymerase and the corresponding reaction buffers. Genotype PCR assembly reactions included a template (20-200 ng gDNA or cDNA), dNTPs at a 200 μM final concentration, forward and reverse primers at a final concentration of 300 nM (for Taq reaction) or 500 nM (for Q5 reaction), a polymerase-specific reaction buffer at a 1x final concentration, and Q5 at 0.02 U μl^−1^ or Taq at 0.025 U μl^−1^. These components were scaled to 25 μl reaction volumes. See **Supplementary Table 1** for primer sequences and cycling conditions. sgRNA templates were prepared as described in [16–18] [44]. Briefly, a universal reverse primer was combined with a forward primer containing a 5’ T7 polymerase binding site, a gene-specific target sequence, and approximately 20 nucleotides of a 3’ sequence complementary to the universal reverser primer in a 100 μl reaction at a 500 nM final concentration for each primer, dNTPs at a 200 μM final concentration, Q5 reaction buffer at a 1x final concentration, and 2U of Q5. PCR products were visualized via agarose gel electrophoresis and nucleic acid staining with SYBR safe DNA stain (S33102, Thermo Fisher Scientific), and imaged using an EZ Gel Documentation System (Bio-Rad, 1708270 and 1708273). The PCR products were purified using the PCR QIAquick PCR cleanup kit (Qiagen, 28106) according to the manufacturer’s instructions.

### RNA isolation

Total RNA was isolated from embryonic or larval tissue by mechanically homogenizing the tissue at room temperature in 200-500 μl TRIzol (Ambion, 15596-018) followed by RNA isolation according to the TRIzol product instructions or using a Direct-zol RNA MiniPrep Plus kit (ZYMO Research Corp, 2072). DNA contamination was removed from the TRIzol-isolated RNA via enzymatic digestion with 10 U of Turbo DNase (Ambion, AM2239) at 37°C for 15 minutes in a reaction tube for TRIzol-mediated extraction or on a ZYMO RNA MiniPrep spin column. DNase was removed from the RNA via organic extraction with phenol:CHCl3:IAA (isoamyl alcohol) (125:24:1) followed by CHCl3:IAA (24:1), then precipitated by adding 10% (v/v) 3M pH 5.2 sodium acetate solution and 2.5 volumes of 100% ice-cold ethanol and cooled to −20°C for ≥20 minutes, then centrifuged at 16,000-20,000 RCF for 20 minutes. The RNA pellet was washed twice with 70% (v/v) EtOH, air-dried, and dissolved in 20-50 μl DNase/RNase-Free water. The RNA isolated using the ZYMO Direct-zol RNA MiniPrep columns was eluted in 50 μl of DNase/RNase-Free water. The final concentrations were measured at a 260 nm/280 nm absorbance on a Nanodrop 2000 spectrophotometer.

### cDNA synthesis for cloning

Up to 1 μg of DNA-free RNA was reverse transcribed using ProtoScript II Reverse Transcriptase (NEB, M0368) and anchored oligo dT primers according to the product instructions.

### T7E1 mutant survey (F0)

T7 endonuclease 1 (#E3321, New England BioLabs) was used to survey for heteroduplexed PCR products as a result of mutagenized target loci. 200 ng of PCR product was denatured and reannealed by heating to 95°C for 5 minutes followed by gradual cooling to 85°C at a rate of 0.5°C/second and then to 25°C at a rate of 0.1°C/second. Annealed DNA was exposed to T7E1 for 15 minutes at 37°C followed immediately by cooling ice. Products were separated and visualized on 2% agarose gel alongside 200 ng of undigested product for comparison.

### Outcross and T7E1 mutant survey (F1)

Sibling larvae (to the injected embryos positive in the T7E1 mutant survey) were raised to sexual maturity and five adult individuals were crossed with wild-type AB adults. Fifteen embryos from each cross were pooled and gDNA was isolated and cleaned as done previously, and dissolved in 100 μl nanopure water. PCR amplification of target loci was done as previously described and products were column purified and eluted in 20 μl Elution Buffer (Qiagen). T7E1 survey was performed as described above. Sibling embryos to T7E1 positive extracts were reared as putative cyp20a1 heterozygotes, whereas those that were T7E1 negative were euthanized.

### Morphological observations

Zebrafish larvae of the AB and *cyp20a1*−/− mutant line were kept until 6 dpf in 35 mm culture dishes (Falcon) containing 10 larvae in 10 mL per dish. At 6 dpf, the larvae were visually compared using a stereomicroscope, scored based on swim bladder inflation, and imaged. This experiment was independently repeated three times with six dishes per line (AB, wh^61^) and experiment (total *n* = 18). The morphological appearances of adult AB and mutant fish were also compared during the novel tank assay.

### Behavioral assays

Optomotor response (OMR) assays were performed in “raceway”-shaped arenas created with 2% (w/v) agarose in deionized water with 60 mg L^−1^ Instant Ocean using a custom plastic mold. This mold was modified from a previously published design [19]. Each mold would cast a 7.5 cm × 11.6 cm gel containing 10 individual 7 cm × 0.8 cm raceways. Especially developed plastic molds measuring 11.7 cm × 7.6 cm × 5 mm were custom-built in-house. The molds were then used to create lanes using agarose poured into single-well plastic plates measuring 12.4 cm × 8.1 cm × 1.2 cm (Thermo Scientific). The molds contained five lanes in which the sides were angled at 60° to facilitate visualization. The lanes in the molds were 3.5 mm high with a base of 18 mm at the top, which tapered to 14 mm at the bottom of the lane. There was a 4 mm gap between the lanes in the mold. The agarose lanes were only used once per experiment and were discarded after each use. Videos of sinewave gratings for entrainment were provided by Dr. Elwood Linney. Prior to the video recordings, individual fish were transferred into each raceway and allowed to acclimate for 5 minutes in lighted conditions. Video recordings were acquired with two Logitech C920 USB webcams at a resolution of 960 × 720 pixels and a frame rate of 30 frames second^−1^ (fps) as described in a previous study [20]. A total of 120 larvae per fish line were recorded before the videos were analyzed using custom R scripts.

Standard light-dark locomotor assays were performed using a DanioVision^™^ observation chamber (Noldus Inc.; Wageningen, Netherlands). At 6 dpf, zebrafish AB and *cyp20a1*−/− mutant larvae (wh^60^ and wh^61^) were randomly distributed in 48-well plates and acclimated in the light for 30 minutes prior to the start of the light-dark transitions. Three 10-minute dark periods were each followed by a 10-minute light period. Each replicate experiment was run at approximately the same time of day (early afternoon). The experiments were repeated at least three times with cohorts from separate breeding events (total *n* = 120 for wh^60;^ total *n* = 72 for wh^61^), and the data from the replicate experiments were pooled for final analysis. Videos were recorded at 30 fps and analyzed with EthoVision XT^®^ 12 (Noldus Inc.).

Vibroacoustic startle latency was assessed as described previously [21, 22] and the same set-up was used to test for startle habituation as a form of non-associative learning in 6 dpf larval zebrafish. For each trial, 16 larvae with inflated swim bladders were distributed in a 4×4 acrylic well-plate which was mounted on a minishaker (Brüel & Kjaer, Vibration Exciter 4810) connected to an amplifier (Brüel & Kjaer, Power Amplifier Type 2718). For the startle response assay, vibro-acoustic stimuli were delivered at four different amplitudes (32, 38, 41, 43 dB) and for each amplitude, the stimulus was delivered four times spaced 20 seconds apart. For the habituation assay, vibro-acoustic stimuli were delivered at 43 dB only. To establish a baseline response in the startle habituation assay, the interval of the first three stimuli was set to 2 minutes. The interval of the following 30 stimuli was set at 10 seconds to test for habituation. After an additional 5 minutes of rest, responsiveness recovery was tested after a single stimulus. The startle response was tracked at 1000 fps using a high-speed video camera (Edgertronic, CA) and analyzed using FLOTE [23] and the analysis pipeline developed by [24]. To assess habituation, the fraction of the 16 larvae per plate and stimulus that responded with a short-latency C-bend (SLC; within 15 ms) was calculated. Both the startle response assay (total *n* = 144 larvae) and the habituation assay (total *n* = 11 plates) were repeated three times with cohorts from separate breeding events.

The novel tank assay assesses anxiety-like behaviors and was performed using adult (10-month-old) zebrafish. The experimental room was heated to 26 °C before the assay, which was performed between 11 am and 3 pm. Two narrow tanks (H: 15.1 cm; L: 21.5 cm; W: 5.1 cm) filled with system water were placed next to each other. For each round, two zebrafish (one fish of each line and of the same sex) were placed individually in a 50 mL glass beaker with 2 mL of system water for 30 seconds prior to releasing the fish simultaneously in the novel tank environment [25]. Videos were recorded with a Sony HD HDR-CX5 for 10 minutes. The temperature of the tank surfaces was regularly checked using an infrared thermometer and kept between 24.2 and 27.2 °C.

DeepLabCut (version 2.2.b8) was used to track the zebrafish in the novel tank assay [26, 27]. To enhance the tracking performance, several body parts were labeled, including the snout, left eye, right eye, left gills, right gills dorsal fin, upper caudal fin, base caudal fin, and lower caudal fin. The residual neural network ResNet-50 was trained using 62 manually labeled frames from 5 randomly selected videos, after which 95% of the frames were used for 100,000 training iterations. We validated the training dataset and found the Root Mean Square Error for test was 20.9 pixels and for train: 2.8 pixels (the image resolution was 1920 by 1080 pixels). We then used a p-cutoff of 0.9 to condition the x,y coordinates for future analysis. Ultimately, the x,y values for the snout generated by DeepLabCut were processed using the NTD analysis script to evaluate ‘Total distance moved’, ‘Time percent in bottom third’, ‘Latency for first entry to upper half’, ‘Latency for second entry to upper half’, ‘Number of transitions to top half’, ‘Number of erratic swimming episodes’, ‘Number of freezing episodes’, and ‘Total freeze time’ [28].

### Statistical analysis

Biological data, in particular behavioral data, exhibit inherently wide sample-to-sample variability, and therefore many samples are required to achieve sufficient statistical power for a reliable *p*-value interpretation [29]. As an alternative to null hypothesis significance testing, which focuses on a dichotomous reject-nonreject decision strategy based on *p* values, estimation statistics report on the estimation of effect sizes (point estimates) and their confidence intervals (precision estimates). In this study, we used estimation statistics and depicted effect size using Gardner-Altman plots [30]. For those unfamiliar with interpreting effect sizes, *p* values from unpaired t-tests (parametric) or Mann-Whitney tests (nonparametric) were also calculated and are reported alongside confidence intervals in the following format: ‘mean difference’ (95% confidence intervals; upper limit, lower limit), *p*-value. Normality was determined using the D’Agostino & Pearson test. The statistical results of all assays are listed in **Table S2**.

## RESULTS

### CYP20A1 mutant lines

*CYP20A1* was simultaneously targeted by two different sgRNAs in the 2^nd^ and 3^rd^ exons, resulting in multiple INDEL mutations (**Figure 1A**). Standard F0 outcrossing and sibling incrossing resulted in stable *cyp20a1*−/− mutant lines in the AB background (**Figure 1B, C**). Two separate *cyp20a1*−/− mutant lines were isolated: line 60 (wh^60^), with a 6 bp deletion and 5 bp insertion in exon 2, and line 61 (wh^61^), with a 1 bp insertion in exon 2, and a 6 bp deletion in exon 3 (**Figure 1D**). In both cases, apparent nonsense mutations were created and computational translation of the mutant alleles showed the predicted amino acid sequence (**Figure 1E**). Due to the unavailability of specific antibodies, we were unable to confirm that the CYP20A1 protein was completely absent from these lines, although without the heme-binding domain any protein would be inactive. The behavior concordance (see below) suggests that both lines are missing active Cyp20a1 protein.

**Figure 1.**
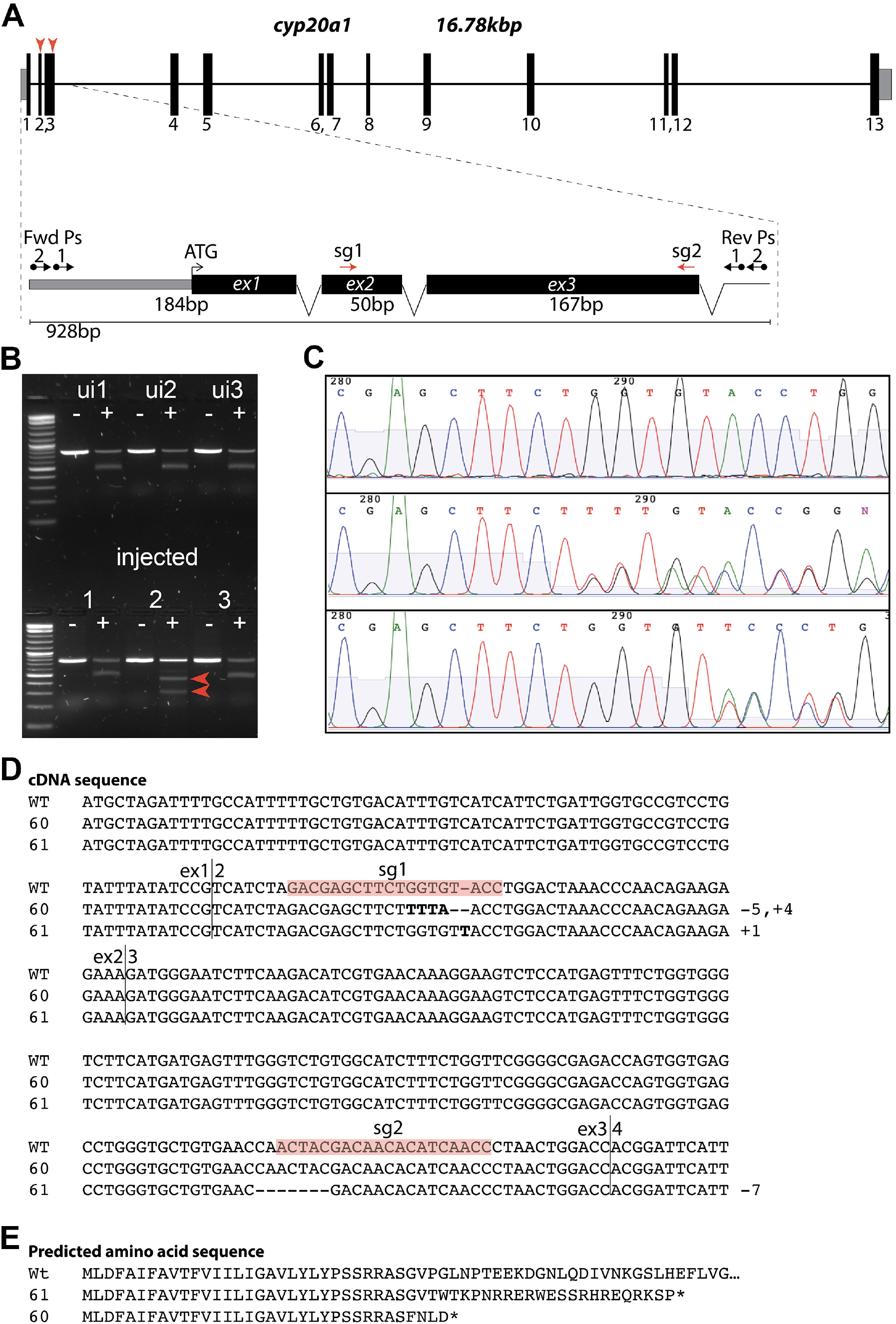
Zebrafish *cyp20a1* gene map and allele sequences. **(A)** Gene models **(B)** Gel image showing PCR products derived from T7E1 mutant survey (F0). Lower gel shows the positive (heteroduplexed) T7D1 signature. **(C)** Chromatogram from F1 Hets (wh60 and wh61) beginning near the sg1 site. (Note the appearance of double peaks). **(E)** cDNA sequences for exons 1-3 for wt, wh^60^, wh^61^. Note wh^60^ is a 5 bp deletion and 4 bp insertion in exon 2 and wh^61^ is a 1 bp insertion in exon 2, and a 6 bp deletion in exon 3. **(D)** Putative translation of cDNAs.

We observed mild morphological differences between the mutant line (wh^61^) and wild-type AB fish in our facility. Fewer mutant fish exhibited swim bladder inflation at 6 dpf than control fish (**Supplementary Figure S1**). Collating the three trials to assess swim bladder inflation showed that the unpaired mean difference of *cyp20a1*−/− wh^61^ (*n* = 18) minus AB (*n* = 18) was −30.6% (95 CI; −38.3, −22.8), *p* < 0.001. In adults, there was a consistent color difference, with the wh^61^ line exhibiting an overall paler pigmentation. (The wh^60^ line was not available to be observed for swim bladder or color at the time this was noted.)

### Larval behavior

We assessed the optomotor response (OMR) of larval zebrafish by analyzing the swimming responses (entrainment) to repeated sinewave gratings moving in one direction and then reversing. The OMR is essential for many animals to correct for deviation from an intended track direction requiring integration of both visual and movement functions. Changes in OMR are indicative of altered motor control, which can originate from altered muscular or retinal sensitivity or neuronal function of the underlying circuit. Wild-type and *cyp20a1*−/− wh^61^ fish were subjected to two instances of OMR visual stimulation (first to the right, then to the left). The *cyp20a1*−/− larvae were less active compared to the wild-type AB strain during the 60 seconds prior to the beginning of the sinewave movement in both the right and left directions (**Figure 2A**). For example, in the 15 seconds before the sinewave moment to the right, the cyp20a1−/− mutant larvae moved on average 0.555 cm less (95CI; −0.841, −0.283), *p* = 0.003. Once the sinewave movement was started, however, both the *cyp20a1*−/− mutant and AB larvae responded equally to the sinewave movement in both directions. **Supplementary Figure S2** indicates the parameters calculated from the larval movement in the 5 minutes prior to the OMR assay. Despite a decrease in average speed by −0.499 mm s^−1^ (95CI; −0.712, −0.287), *p* = 0.001 (**Figure S2A**), distance traveled by −149 mm 5 min^−1^ (95CI; −213, −85.6), *p* = 0.001 (**Figure S2B**), and overall activity prior to the sinewave movement by −14.1% (95CI; −21.3, −6.88), *p* = 0.001 (**Figure S2C**) in *cyp20a1*−/− mutant larvae compared to the AB strain, both AB and *cyp20a1*−/− exhibited a capacity to engage equally in high-speed swimming activity after the sinewave movement was initiated (**Figure S2D**). Collectively, these observations reveal that *cyp20a1*−/− fish are far more reactive to the OMR visual stimulus, despite being less active in the absence of it in this assay.

**Figure 2.**
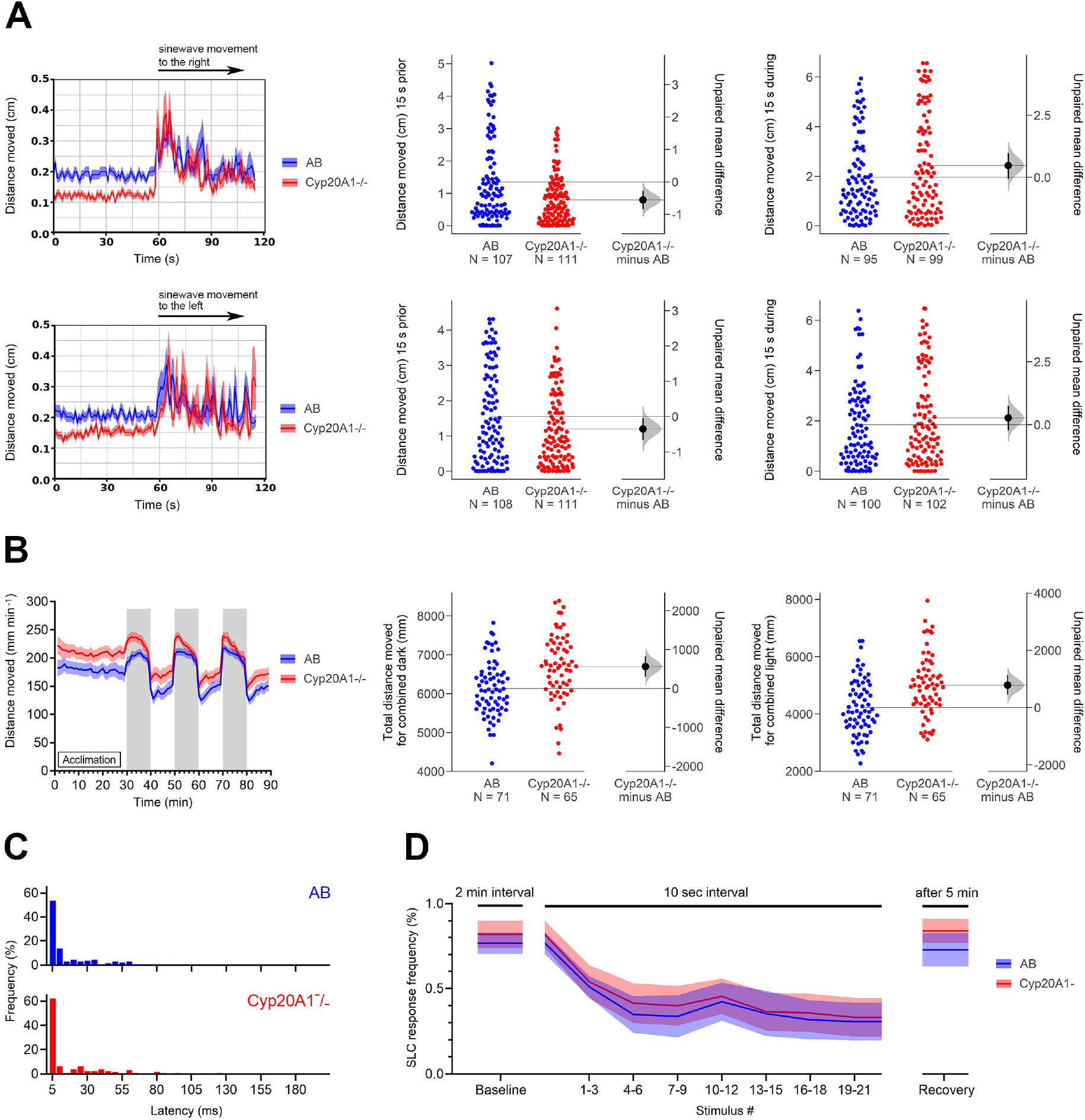
Larval behavior. (**A**) Optomotor response of AB and CYP20A1−/− wh^61^ mutant larvae (*n* = 120). (**B**) The locomotor activity of CYP20A1−/− wh^61^ mutant larvae (*n* = 65) during the dark and the light phases in comparison to the AB larvae (*n* = 71). (**C**) Rapid startle response to the highest acoustic stimulus (43 dB) of AB (*n* = 138) and CYP20A1−/− wh^61^ (*n* = 127) larvae. (**D**) Habituation to the highest acoustic stimulus measured as short-latency C-bend response (< 15 ms) per plate (*n* = 11) and depicted as mean ± 95 CI. All individual data points represent biologically independent replicates from three independent experiments.

Larval locomotion during daylight in some fish species is driven by a natural need for hunting and exploring. Upon sudden darkness, zebrafish larvae respond with hyperactivity, potentially in response to an overshadowing predator. We used a light-dark assay consisting of a 30-minute light acclimation period followed by repeating 10-minute dark and light exposures. Compared to the AB strain, the locomotor activity in *cyp20a1*−/− wh^60^ and wh^61^ mutant larvae was higher during the acclimation period, as well as during the dark stimulations (both wh^60^ and wh^61^ at *p* < .0001), whereas the wh^61^ mutants also exhibited hyperactivity in the light phases following the dark stimulations (*p* < .0001, **Figure 2B, Supplementary Figure S3**). The difference in response of wh^60^ and wh^61^ mutant larvae suggests that in one of the mutants some residual gene product is being produced, contributing to the difference in response during the light phase. Both mutants exhibit hyperactivity, suggesting that *cyp20a1*−/− behavioral differences may not be attributed to muscle impairments but rather to neurological or other effects. This is further supported by the fact that the *cyp20a1*−/− fish remained less active than the AB fish during the OMR assays just prior to any visual stimulus (**Figure 2A**), but significantly increased their locomotor activity during the first 15 seconds of the OMR stimulus.

The startle response in fish is triggered by sensory stimuli (visual or vibro-acoustic) to rapidly escape from predators and changes in this response can be indicative of altered neuronal cell development or transmission. The startle latency exhibited by the *cyp20a1*−/− wh^61^ mutant larvae did not differ from that of the AB larvae at the highest three stimulus intensities (**Figure 2C, Supplementary Figures S4**) but showed on average a more rapid response at the two lowest stimulus intensity (*p* < .017, **Supplementary Figures S4**). It is common for some larvae to not exhibit a startle response when exposed to a vibro-acoustic stimulus. However, AB larvae were more responsive than *cyp20a1*−/− wh^61^ mutant larvae at the lowest and highest stimulus intensity (*p* < .002, *p* < .008, **Supplementary Figures S4C**). Repeated stimulation within a short period often leads to habituation, indicating that the nervous system is capable of filtering out irrelevant information. However, this can be impaired in several psychiatric and neurological diseases including schizophrenia and autism. Both the AB larvae and the *cyp20a1*−/− wh^61^ mutant larvae appeared to adapt to the highest auditory stimulus, suggesting habituation (**Figure 2D**).

### Adult behavior

We also examined adult behaviors that are related to anxiety disorders using the novel tank assay [31]. This behavioral assay involves an anxiety response to a novel environment, and by repeating the assay a measure of acclimation or, conversely, a buildup of stress can also be achieved. In all three trials, the *cyp20a1*−/− fish (wh^61^) spent more time in the bottom third of the novel tank (**Figure 3A**), which is an indication of anxiety-like behavior. Both the *cyp20a1*−/− and the AB fish showed a tendency toward increased bottom-dwelling when the assay was repeated on days 7 and 14, suggesting a long-lasting stress effect from the handling in the previous week. Consistent with the increased time spent in the bottom third of the tank, the *cyp20a1*−/− fish also showed a delay in moving to the top half of the tank, after the first (**Supplementary Figure S5A**) and the second entry (**Supplementary Figure S5B**). There also was a decreased number of transitions to the top half (**Supplementary Figure S5C**). For statistical results of all three trials please see **Table S2**.

**Figure 3.**
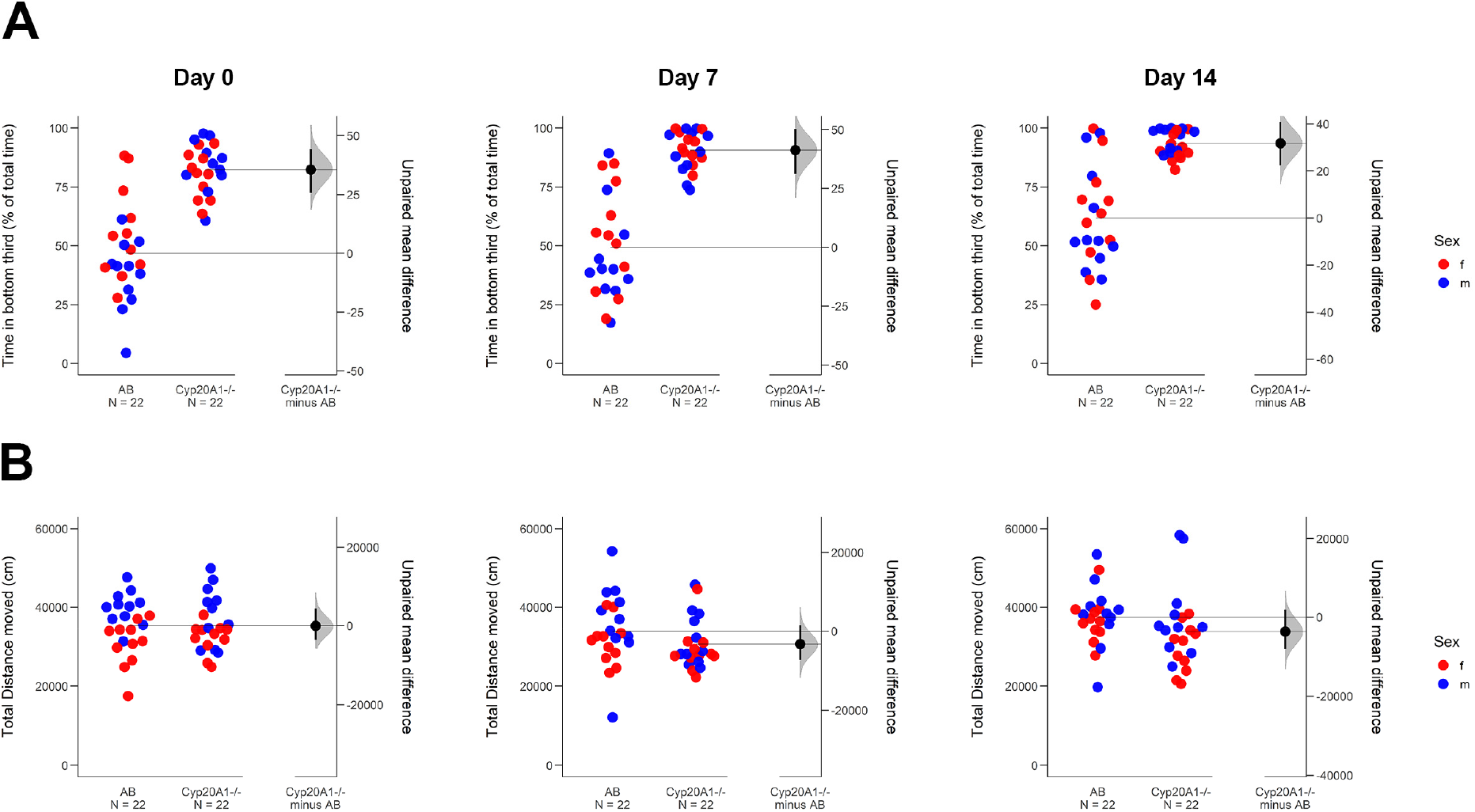
Adult behavior in the novel tank assay. (**A**) CYP20A1−/− wh^61^ mutant zebrafish spend more time in the bottom third of the novel tank in the first 10 minutes in comparison to AB zebrafish. (**B**) The distance moved in the novel tank does not differ between CYP20A1−/− wh^61^ mutant zebrafish and AB zebrafish. The experiment was repeated on day 7 and day 14 with the same fish.

In terms of distance moved, both the *cyp20a1*−/− and the wild-type AB fish moved about the same (**Figure 3B**). The total duration of time spent freezing (displacement of ≤ 3 mm/s, **Supplementary Figure S5D**), the number of freezing episodes (at least 1 s of immobility, **Supplementary Figure S5E**), and the number of erratic swimming movements (darting, **Supplementary Figure S5F**) differed based on the *p*-value in one out of three trials between the *cyp20a1*−/− and the wild-type AB zebrafish. Estimation statistics only indicated a decrease in the number of erratic swimming movements in *cyp20a1*−/− fish. No difference between males and females was observed for the endpoints measured except for the distance traveled in the first trial, in which both *cyp20a1*−/− and the AB females generally moved less than males.

The novel tank assay was performed using two tanks to record a *cyp20a1*−/− fish and an AB fish of the same sex at the same time. This setup allowed for direct visual comparison of the adult morphology, which in every case indicated a paler appearance of the *cyp20a1*−/− zebrafish in comparison to the AB.

## DISCUSSION

Our specific focus on behavior was prompted by the possible neurological implications of *CYP20A1* RNA expression levels in the hippocampus and substantia nigra in the human brain [5], early larval zebrafish, [11], and in the developing mouse brain [13]. Moreover, our prior studies with transient morpholino knockdown of *cyp20a1* resulted in behavioral phenotypes involving visual responses and overall activity, akin to hyperactivity [6]. The results from our CRISPR/Cas9 *cyp20a1*−/− mutant experiments further support the idea that the function(s) of CYP20A1 are involved in neurological processes that when disrupted lead to behavioral changes in zebrafish.

In an earlier study [6], we gleaned information from case reports of interstitial micro-deletions in the human Chr2q33.1-2q33.2 region, including *CYP20A1* gene loss, which resulted in a suite of neurological defects among other adverse effects [6]. Patients with 2q33 microdeletion syndrome display developmental delays, psychomotor retardation, hyperactivity and bouts of anxiety, and in some cases delayed visuomotor coordination [32, 33]. However, hyperactivity, particularly in children, was observed primarily in patients in which the deletions in this region included the locus for *CYP20A1*. Recent examination of additional case studies [34] has now strengthened this observation of possible involvement in human neurobehavioral disorders.

Zebrafish inherently exhibit many different types of behavior, some of which are analogous to mammalian behaviors. These include anxiety and hyperactivity [31, 35, 36]. These cross-species behavior analogies are cemented by the observations of identical outcomes resulting from pharmacological manipulations. For instance, ethanol reduces stress and anxiety behaviors, resulting in increased exploration and reduced erratic movements, whereas caffeine increases stress-associated behaviors, resulting in irregular movements [37]. Such observations often are in parallel with shifts in cortisol levels, which are used as a physiological marker of anxiety and stress [38, 39]. In our study, the dark-induced hyperactivity in *cyp20a1*−/− mutant larvae, and the finding that *cyp20a1*−/− adults spent more time in the bottom third of a novel tank compared to wild-type fish, suggest that the absence of *Cyp20a1* gene product may dysregulate steroid hormones such as cortisol. The resemblance in anxiety and hyperactivity responses between humans and zebrafish with deletions in the *Cyp20a1* locus suggests that our zebrafish *cyp20a1*−/− crispants can serve as a disease model organism.

The endogenous catalytic function of CYP20A1 remains unknown. Earlier, based on predicted protein structural features, we speculated that substrates of CYP20A1 may carry their own oxygen for catalysis and that these might include oxysterols or related compounds [6]. Human *cyp20a1* expressed in yeast has been reported to weakly act on non-physiological luminogenic substrates and can be inhibited by azoles, suggesting that this enzyme may catalyze typical P450-type transformations, albeit at low reaction rates [8]. However, this observation may aid in our search to determine whether candidate biological substrates are detectably metabolized by recombinant proteins.

Although the catalytic function of CYP20A1 remains elusive, its broad tissue distribution suggests that CYP20A1 likely possesses multiple catalytic activities. Additionally, the primary activity of a given protein is broadly affected by common processes in different organs, including the brain. CYP20A1 is widely distributed in the animal kingdom, including in early-diverging groups such as sponges [40]. Although CYP20A1 appears to be ubiquitous among deuterostomes, its presence is sparse among arthropods [40], apparently having been lost in some groups. Nevertheless, the broad distribution suggests that this orphan P450 may serve functions that are critical in vertebrate biochemistry and that these may be conserved among animals, especially in the deuterostome lineage.

As with function, the regulation of CYP20A1 expression is not understood. Most human and macaque tissues exhibit some level of expression at the RNA level [5, 41]. We also found *CYP20A1* expression in most tissues of adult zebrafish [6], and widespread expression has been found in mice [13]. Unusual among non-mitochondrial P450s, the N-termini of the predicted CYP20A1 protein sequences are nearly identical across mammals [6], suggesting a conserved targeting or functioning of this protein region.

Although we believe that CYP20A1 has role(s) in neural tissues, the expression patterns clearly imply functions in other tissues. Tissue expression and promoter analysis also suggest reproductive, immune, hematopoietic, and neural involvement. Previously, we reported that *cyp20a1* transcript expression in zebrafish embryos is modestly affected by steroids and other nuclear receptor agonists, and was suppressed by the neurotoxicant methylmercury [6]. In any case, the behavioral alterations in zebrafish in which *cyp20a1* has been knocked down [6] or knocked out (*cyp20a1*−/−; this study) imply that if there are multiple functions for this protein, these must include function(s) in the brain and steroid hormone synthesizing gonads.

The expression of *CYP20A1* transcript during development and in multiple adult organs in mammals and zebrafish implies endogenous regulation. In a human tissue screen, the highest levels of CYP20A1 expression were observed in endocrine tissues (as a group) and the pancreas [5]. We previously found the highest expression level in adult zebrafish gonads [6]. Multiple other lines of experimental evidence point to endocrine participation involving steroids, which is consistent with the expression patterns in fish and humans. The hyperactivity in larvae and the anxiety-like behavior in adults may indicate a dysregulation of glucocorticoid biochemistry as previously described for these specific behaviors [42, 43]

In summary, we report on a *cyp20a1*(−/−) crispant zebrafish and the results obtained substantiate the specific involvement of Cyp20a1 in behavioral phenotypes in this vertebrate model. However, the broader significance of Cyp20a1 to vertebrate physiology and disease processes remains unclear. The fact that the *cyp20a1*−/− null strain grows and reproduces with few defects suggests that *cyp20a1* is not an essential gene, barring some escape from the mutant condition or low-level redundancy as seen with some other genes, including in zebrafish (e.g., [44]). A comprehensive search for substrates is underway with recombinant zebrafish Cyp20a1 expressed in *E. coli*, and metabolomics studies. The features of CYP20A1 structure, regulation, and biological correlations aid in the deciphering of the molecular functions and roles of this orphan P450 in health and disease, as well as the evolution of these functions. The mutant strains we have developed are being explored to determine the functional and metabolic significance of CYP20A1. The CRISPR/Cas generated *cyp20a1*−/− zebrafish described herein will enable the functional characterization of this last human orphan P450, potentially advancing our understanding of the molecular mechanisms related to human mental health and the search for potential therapies.

## Acknowledgments

These studies were supported in part by the Boston University Superfund Research Program NIH 5P42ES007381 (MCS, NRB, FXM, JVG, JJS), the Woods Hole Center for Oceans and Human Health (NIH: P01ES021923 and P01ES028938; NSF: OCE-1314642 and OCE-1840381; NRB and JJS), and EBI/EMBL Medakatox NIEHS R01ES029917 (JVG). DCL was funded by a UK-US Fulbright Scholarship.

