## Supplemental Material for "Orphan cytochrome P450 20A1 CRISPR/Cas 9 mutants and neurobehavioral phenotypes in zebrafish"

**Table S1**. Primer sequences for endpoint PCR and CRISPR-Cas sgRNA target and synthesis sequences.

| **Sequence Name** | **Barcode ID** | **Primer sequence (5')** | **Cycling condition** | **Polymerase** | **Product length (bp)** |
|---|---|---|---|---|---|
| cyp20-wtP2-F | 1025475438 | TGTCATCATTCTGATTGGTGCC | 98˚ - 3 min - (1x); 98˚ - 10 sec, 65˚ - 10 sec, 72˚ - 20 sec (35x); 72˚ - 5 min (1x) | NEB Q5 | 437 |
| cyp20-wtP2-R | 1025479906 | GTTGATGTGTTGTCGTAGTTGG | | | |
| cyp20-ms61P1-F | 1025475440 | TAGACGAGCTTCTGGTGTTACC | 98˚ - 3 min - (1x); 98˚ - 10 sec, 65˚ - 10 sec, 72˚ - 20 sec (35x); 72˚ - 5 min (1x) | NEB Q5 | 277 |
| cyp20-ms61P1-R | 1025475441 | CAGTTAGGGTTGATGTGTTGTCG | | | |
| cyp20_60_wtF | 1033095856 | GAC GAG CTT CTG GTG TAC CTG G | 98˚ - 3 min - (1x); 98˚ - 10 sec, 68˚ - 10 sec, 72˚ - 20 sec (35x); 72˚ - 5 min (1x) | NEB Q5 | 304 |
| cyp20_60_wtR | 1033095857 | CAC TGC GGC TAC TCA CTG GT | | | |
| cyp20_60_mutF | 1033095860 | TTA CGC CCC TGT CTT GCA GT | 98˚ - 3 min - (1x); 98˚ - 10 sec, 65˚ - 10 sec, 72˚ - 30 sec (35x); 72˚ - 5 min (1x) | NEB Q5 | 744 |
| cyp20_60_mutR | 1033095861 | CTG TTG GGT TTA GTC CAG GTT AAA AG | | | |
| dr.cyp20a1-F | 1023898079 | ACCATGCTAGATTTTGCCATTTTTGCTGTG | | | |
| dr.cyp20a1-R | 1023898080 | TCAGTTTCTCTTGCTGACCGTG | | | |
| z.cyp20a1-5'utr-F | 1023898081 | GTAGTCGAGTACCGATCTAGAGG | | | |
| z.cyp20a1-3'utr-R | 1023898082 | GTGTAATTCCCATCCTCCAGAGG | | | |
| dr.cyp20a1.T7.sg1 | 1023342952 | GATTAATACGACTCACTATAGGACGAGCTTCTGG TGTACCGTTTTAGAGCTAGAAATAGC | | | |
| dr.cyp20a1.T7.sg2 | 1023342953 | GATTAATACGACTCACTATAGGTTGATGTGTTGTC GTAGTGTTTTAGAGCTAGAAATAGC | | | |
| sgRNA Universal reverse primer | | AAAAGCACCGACTCGGTGCCACTTTTTCAAGTTGATAACGGACTAGCCTTATTTTAACTTGCTATTTCTAGCTCTAAAAC | | | |

| | | |
|---|---|---|
| dr.cyp20a1.seqpF1 | 1023342954 | ATCGCCAGCTCGTAGTTCAC |
| dr.cyp20a1.seqpR1 | 1023342955 | CAGTCTTCAACTGTAAATGCAGC |
| dr.cyp20a1.seqpF2 | 1023342956 | TCCTGATGGTCATTGTAGACG |
| dr.cyp20a1.seqpR2 | 1023342957 | CAGGCGGACTGATAATTCAGG |
| CYP20A1_Dr_pENTRF | 1018501550 | CACCATGCTAGATTTTGCCA |
| CYP20A1_Dr_pENTRR | 1018501551 | TCTCTTGCTGACCGTGATCCA |
| zf_cyp20_f4 | 1017216446 | TACAGGAGGTGGAAGGAAAGGTG |
| zf_cyp20_r4 | 1017216447 | GACGACACCAAGGGCATAGATAAC |

**Table S2**. Statistical results.

| **Assay** | **Endpoint** | **CYP20A1 Line** | **# of Trials** | **Total *n* (AB)** | **Total *n* (Cyp20A1)** | **Estimation Stats** | **Passed Normality Test** | **Unpaired t-test[1]** | **Mann-Whitney Test[2]** |
|---|---|---|---|---|---|---|---|---|---|
| Morphology | Swim Bladder Inflation | $wh^{61}$ | 3 | 18 | 18 | Unpaired mean difference of Cyp20A1-/- (n = 18) minus AB (*n* = 18) -30.6 [95CI -38.3; -22.8] | Some | | U = 6, *p* < .001*** |
| OMR | Right Grating (prior) | $wh^{61}$ | 3 | 107 | 111 | Unpaired mean difference of Cyp20A1-/- (*n* = 111) minus AB (*n* = 107) -0.555 [95CI -0.841; -0.283] | No | | U = 4577, *p* = .003** |
| OMR | Right Grating (during) | $wh^{61}$ | 3 | 95 | 99 | Unpaired mean difference of Cyp20A1-/- (*n* = 99) minus AB (*n* = 95) 0.472 [95CI -0.0656; 0.973] | No | | U = 4189, *p* = .190 |
| OMR | Left Grating (prior) | $wh^{61}$ | 3 | 108 | 111 | Unpaired mean difference of Cyp20A1-/- (*n* = 111) minus AB (*n* = 108) -0.349 [95CI -0.668; -0.0377] | No | | U = 5266, *p* = .121 |

| | | | | | | | | | |
|---|---|---|---|---|---|---|---|---|---|
| OMR | Left Grating (during) | wh$^{61}$ | 3 | 100 | 102 | Unpaired mean difference of Cyp20A1-/- (*n* = 102) minus AB (*n* = 100) 0.267 [95CI -0.223; 0.753] | No | | U = 4760, *p* = .414 |
| OMR | Average Speed | wh$^{61}$ | 3 | 115 | 115 | Unpaired mean difference of Cyp20A1-/- (*n* = 115) minus AB (*n* = 115) -0.499 [95CI -0.712; -0.287] | Yes | t(228) = 4.711, *p* < .001*** | |
| OMR | Maximum Speed | wh$^{61}$ | 3 | 115 | 115 | Unpaired mean difference of Cyp20A1-/- (*n* = 115) minus AB (*n* = 115) 1.11 [95CI -0.293; 3.13] | Some | | U =5778, *p* = .098 |
| OMR | Activity | wh$^{61}$ | 3 | 115 | 115 | Unpaired mean difference of Cyp20A1-/- (*n* = 115) minus AB (*n* = 115) -14.1 [95CI -21.3; -6.88] | No | | U = 4492, *p* < .001*** |
| OMR | Total Distance Traveled | wh$^{61}$ | 3 | 115 | 115 | Unpaired mean difference of Cyp20A1-/- (*n* = 115) minus AB (*n* = 115) -149 [95CI -213; -85.6] | Yes | t(228) = 4.706, *p* < .001*** | |
| Ligth-Dark | Total Activity Dark | wh$^{61}$ | 3 | 71 | 65 | Unpaired mean difference of Cyp20A1-/- (*n* = 65) minus AB (*n* = 71) 566 [95CI 299; 824] | Yes | t(134) = 4.251, *p* < .0001**** | |
| Ligth-Dark | Total Activity Light | wh$^{61}$ | 3 | 71 | 65 | Unpaired mean difference of Cyp20A1-/- (*n* = 65) minus AB (*n* = 71) 782 [95CI 448; 1130] | Yes | t(134) = 4.447, *p* < .0001**** | |
| Ligth-Dark | Total Activity Dark | wh$^{61}$ | 5 | 120 | 120 | Unpaired mean difference of Cyp20A1-/- (*n* = 120) minus AB (*n* = 120) 659 [95CI 433; 883] | Some | | U = 4200, *p* < .0001**** |
| Ligth-Dark | Total Activity Light | wh$^{61}$ | 5 | 120 | 120 | Unpaired mean difference of Cyp20A1-/- (*n* = 120) minus AB (*n* = 120) 58.9 [95CI -213; 312] | Yes | t(238) = 0.44399, *p* = .6604 | |
| Startle Response | Startle Latency 32dB | wh$^{61}$ | 3 | 111 | 83 | Unpaired mean difference of Cyp20A1-/- (*n* = 83) minus AB (*n* = 111) -3.34 [95CI -14.1; 8.58] | No | | U = 3685, *p* = .017* |
| Startle Response | Startle Latency 38dB | wh61 | 3 | 133 | 123 | Unpaired mean difference of Cyp20A1-/- (*n* = 123) minus AB (*n* = 133) 2.02 [95CI -5.13; 8.93] | No | | U = 7395, *p* = .185 |
| Startle Response | Startle Latency 41dB | wh61 | 3 | 135 | 124 | Unpaired mean difference of Cyp20A1-/- (*n* = 124) minus AB (*n* = 135) 3.02 [95CI -1.96; 8.68] | No | | U = 7805, *p* = .348 |
| Startle Response | Startle Latency 43dB | wh61 | 3 | 138 | 127 | Unpaired mean difference of Cyp20A1-/- (*n* = 127) minus AB (*n* = 138) 0.0176 [95CI -4.88; 5.28] | No | | U = 7977, *p* = .207 |
| Startle Response | Short Latency C-Bend Bias 32dB | wh61 | 3 | 111 | 83 | Unpaired mean difference of Cyp20A1-/- (*n* = 83) minus AB (*n* = 111) 0.344 [95CI 0.0934; 0.582] | | | |
| Startle Response | Short Latency C-Bend Bias 38dB | wh$^{61}$ | 3 | 133 | 123 | Unpaired mean difference of Cyp20A1-/- (*n* = 123) minus AB (*n* = 133) 0.137 [95CI -0.0474; 0.34] | | | |
| Startle Response | Short Latency C-Bend Bias 41dB | wh$^{61}$ | 3 | 135 | 124 | Unpaired mean difference of Cyp20A1-/- (*n* = 124) minus AB (*n* = 135) 0.0345 [95CI -0.131; 0.212] | | | |

| | | | | | | | | | |
|---|---|---|---|---|---|---|---|---|---|
| Startle Response | Short Latency C-Bend Bias 43dB | $wh^{61}$ | 3 | 138 | 127 | Unpaired mean difference of Cyp20A1-/- (*n* = 127) minus AB (*n* = 138) 0.0327 [95CI -0.141; 0.201] | | | |
| Startle Response | Fraction Responding 32dB | $wh^{61}$ | 3 | 140 | 129 | Unpaired mean difference of Cyp20A1-/- (*n* = 129) minus AB (*n* = 140) -0.148 [95CI -0.24; -0.055] | No | | U = 7092, *p* = .002** |
| Startle Response | Fraction Responding 38dB | $wh^{61}$ | 3 | 140 | 131 | Unpaired mean difference of Cyp20A1-/- (*n* = 131) minus AB (*n* = 140) -0.0154 [95CI -0.0901; 0.0587] | No | | U = 8913, *p* = .637 |
| Startle Response | Fraction Responding 41dB | $wh^{61}$ | 3 | 140 | 129 | Unpaired mean difference of Cyp20A1-/- (*n* = 129) minus AB (*n* = 140) -0.0444 [95CI -0.101; 0.00953] | No | | U = 8122, *p* = .036* |
| Startle Response | Fraction Responding 43dB | $wh^{61}$ | 3 | 140 | 130 | Unpaired mean difference of Cyp20A1-/- (*n* = 130) minus AB (*n* = 140) -0.0536 [95CI -0.103; -0.00462] | No | | U = 8002, *p* = .008** |
| Novel Tank Assay (Trial1) | Time In Bottom Third | $wh^{61}$ | 1 | 22 | 22 | Unpaired mean difference of Cyp20 (*n* = 22) minus AB (*n* = 22) 35.6 [95CI 25.7; 44.4] | Yes | t(42) = 7.423, *p* < .001*** | |
| Novel Tank Assay (Trial2) | Time In Bottom Third | $wh^{61}$ | 1 | 22 | 22 | Unpaired mean difference of Cyp20 (*n* = 22) minus AB (*n* = 22) 41.3 [95CI 31.2; 50.3] | Yes | t(42) = 8.430, *p* < .001*** | |
| Novel Tank Assay (Trial3) | Time In Bottom Third | $wh^{61}$ | 1 | 22 | 22 | Unpaired mean difference of Cyp20 (*n* = 22) minus AB (*n* = 22) 31.8 [95CI 22.3; 40.8] | Yes | t(42) = 6.646, *p* < .001*** | |
| Novel Tank Assay (Trial1) | Latency For First Entry | $wh^{61}$ | 1 | 22 | 22 | Unpaired mean difference of Cyp20 (*n* = 22) minus AB (*n* = 22) 118 [95CI 73.1; 183] | Some | | U = 54, *p* < .001*** |
| Novel Tank Assay (Trial2) | Latency For First Entry | $wh^{61}$ | 1 | 22 | 22 | Unpaired mean difference of Cyp20 (*n* = 22) minus AB (*n* = 22) 151 [95CI 80; 225] | Some | | U = 103.5, *p* < .001*** |
| Novel Tank Assay (Trial3) | Latency For First Entry | $wh^{61}$ | 1 | 22 | 22 | Unpaired mean difference of Cyp20 (*n* = 22) minus AB (*n* = 22) 69.9 [95CI -8.18; 142] | Some | | U = 176, *p* = .123 |
| Novel Tank Assay (Trial1) | Latency For Second Entry | $wh^{61}$ | 1 | 22 | 22 | Unpaired mean difference of Cyp20 (*n* = 22) minus AB (*n* = 22) 128 [95CI 75.1; 193] | Some | | U = 72.5, *p* < .001*** |
| Novel Tank Assay (Trial2) | Latency For Second Entry | $wh^{61}$ | 1 | 22 | 22 | Unpaired mean difference of Cyp20 (*n* = 22) minus AB (*n* = 22) 76.6 [95CI -8.86; 161] | Some | | U = 178, *p* = .135 |
| Novel Tank Assay (Trial3) | Latency For Second Entry | $wh^{61}$ | 1 | 22 | 22 | Unpaired mean difference of Cyp20 (*n* = 22) minus AB (*n* = 22) 107 [95CI 9.49; 192] | Some | | U = 159, *p* = .051 |
| Novel Tank Assay (Trial1) | Total Distance Traveled | $wh^{61}$ | 1 | 22 | 22 | Unpaired mean difference of Cyp20 (*n* = 22) minus AB (*n* = 22) -52.6 [95CI -3600; 4390] | Yes | t(42) = 0.02575, *p* = .980 | |
| Novel Tank Assay (Trial2) | Total Distance Traveled | $wh^{61}$ | 1 | 22 | 22 | Unpaired mean difference of Cyp20 (*n* = 22) minus AB (*n* = 22) -3260 [95CI -7310; 1510] | Some | | U = 158, *p* = .049* |

| | | | | | | | | | |
|---|---|---|---|---|---|---|---|---|---|
| Novel Tank Assay (Trial3) | Total Distance Traveled | $wh^{61}$ | 1 | 22 | 22 | Unpaired mean difference of Cyp20 (*n* = 22) minus AB (*n* = 22) -3650 [95CI -8090; 1830] | Some | | U = 147, *p* = .025* |
| Novel Tank Assay (Trial1) | No Transitions Top Half | $wh^{61}$ | 1 | 22 | 22 | Unpaired mean difference of Cyp20 (*n* = 22) minus AB (*n* = 22) -17.2 [95CI -30; -5.28] | Yes | t(42) = 2.595, *p* = .013* | |
| Novel Tank Assay (Trial2) | No Transitions Top Half | $wh^{61}$ | 1 | 22 | 22 | Unpaired mean difference of Cyp20 (*n* = 22) minus AB (*n* = 22) -26.1 [95CI -38.6; -12.9] | Some | | U = 81, *p* < .001*** |
| Novel Tank Assay (Trial3) | No Transitions Top Half | $wh^{61}$ | 1 | 22 | 22 | Unpaired mean difference of Cyp20 (*n* = 22) minus AB (*n* = 22) -26.9 [95CI -37.8; -17] | Some | | U = 74.50, *p* < .001*** |
| Novel Tank Assay (Trial1) | Total Freeze Time | $wh^{61}$ | 1 | 22 | 22 | Unpaired mean difference of Cyp20 (*n* = 22) minus AB (*n* = 22) 1.27 [95CI -5.32; 13.9] | No | | U = 218.5, *p* = .588 |
| Novel Tank Assay (Trial2) | Total Freeze Time | $wh^{61}$ | 1 | 22 | 22 | Unpaired mean difference of Cyp20 (*n* = 22) minus AB (*n* = 22) -0.182 [95CI -9.91; 24.7] | No | | U = 123, *p* = .004** |
| Novel Tank Assay (Trial3) | Total Freeze Time | $wh^{61}$ | 1 | 22 | 22 | Unpaired mean difference of Cyp20 (*n* = 22) minus AB (*n* = 22) -1.32 [95CI -17.5; 16.2] | No | | U = 197, *p* = .297 |
| Novel Tank Assay (Trial1) | No Freezing Episodes | $wh^{61}$ | 1 | 22 | 22 | Unpaired mean difference of Cyp20 (*n* = 22) minus AB (*n* = 22) 1.36 [95CI -4.64; 9.41] | No | | U = 241.5, *p* = .995 |
| Novel Tank Assay (Trial2) | No Freezing Episodes | $wh^{61}$ | 1 | 22 | 22 | Unpaired mean difference of Cyp20 (*n* = 22) minus AB (*n* = 22) -4.23 [95CI -12.1; 6.06] | No | | U = 155.5, *p* = .042* |
| Novel Tank Assay (Trial3) | No Freezing Episodes | $wh^{61}$ | 1 | 22 | 22 | Unpaired mean difference of Cyp20 (*n* = 22) minus AB (*n* = 22) -4.45 [95CI -13.8; 3.23] | Some | | U = 217, *p* = .564 |
| Novel Tank Assay (Trial1) | No Darting Episodes | $wh^{61}$ | 1 | 22 | 22 | Unpaired mean difference of Cyp20 (*n* = 22) minus AB (*n* = 22) -7.68 [95CI -14.8; -1.93] | Yes | t(42) = 2.294, *p* = .027* | |
| Novel Tank Assay (Trial2) | No Darting Episodes | $wh^{61}$ | 1 | 22 | 22 | Unpaired mean difference of Cyp20 (*n* = 22) minus AB (*n* = 22) -6.68 [95CI -13.1; 0] | Yes | t(42) = 1.930, *p* = .060 | |
| Novel Tank Assay (Trial3) | No Darting Episodes | $wh^{61}$ | 1 | 22 | 22 | Unpaired mean difference of Cyp20 (*n* = 22) minus AB (*n* = 22) -5.82 [95CI -12.8; 0.853] | Yes | t(42) = 1.643, *p* = .108 | |

[1] t(degress of freedom) = the t statistic, *p* = p-value

[2] U = the U statistic, *p* = p value

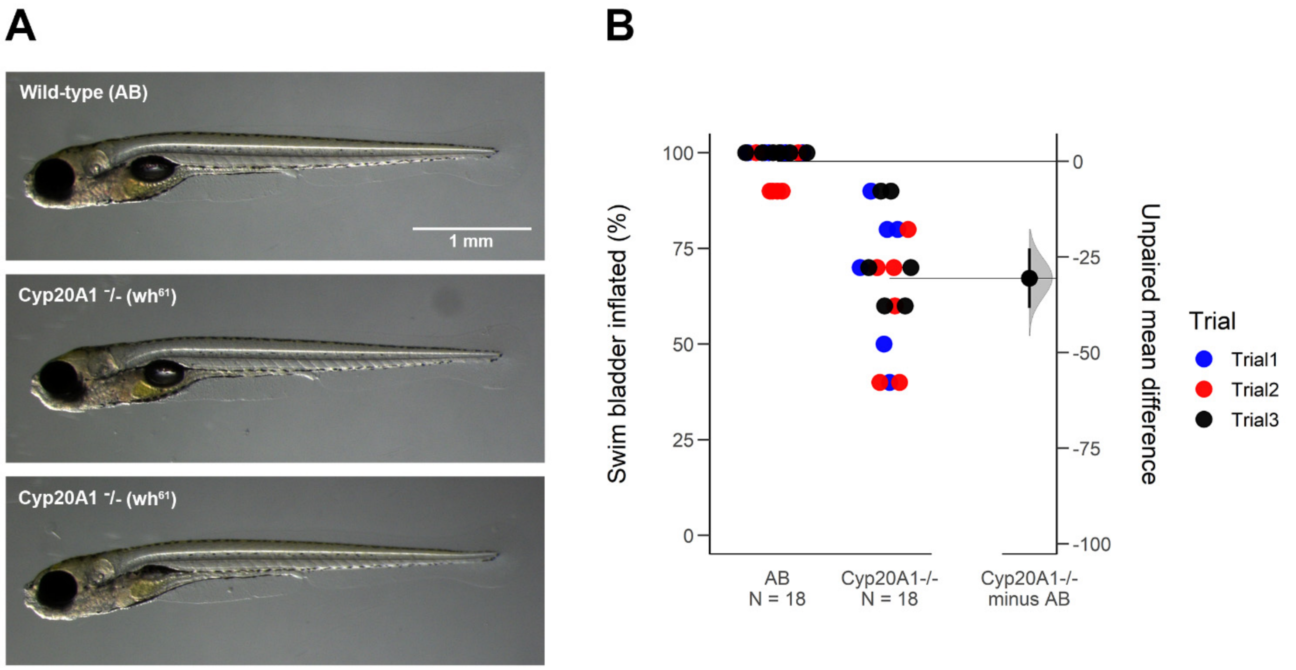


**Figure S1. Morphology of CYP201A1-/- larvae at 6 dpf.** (**A**) CYP201A1 (wh$^{61}$) mutant larvae show no apparent morphological differences in comparison to the wild-type (AB) line except for (**B**) swim bladder inflation, which was reduced in the mutant larvae. The experiment was repeated three times independently with 6 dishes containing 10 larvae per experiment (total *n* of 3 experiments = 18).

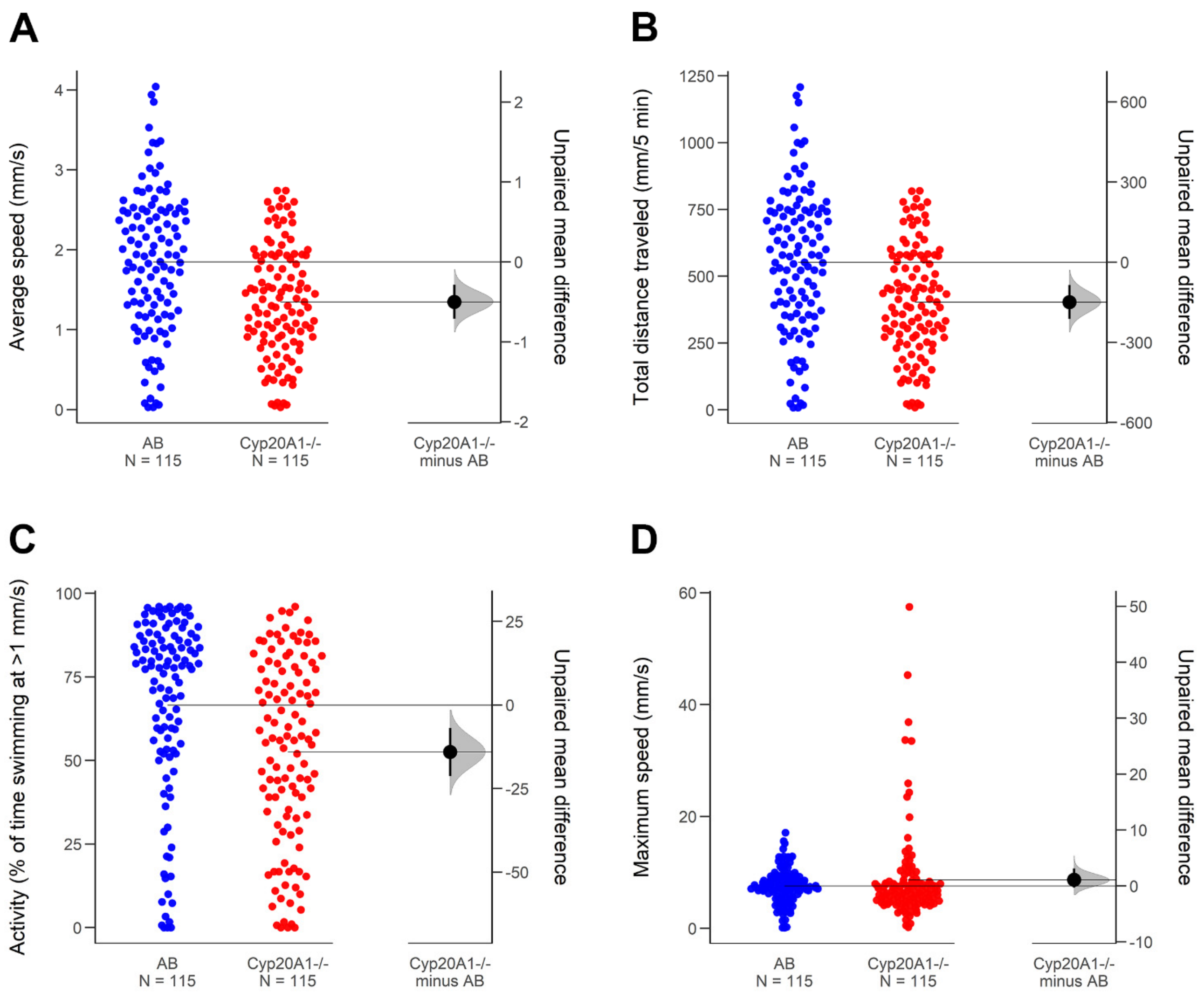


**Figure S2. Activity of larvae prior to optomotor response (OMR) assay.** (**A**) Average speed, (**B**) total distance traveled, (**C**) activity, and (**D**) maximum speed was measured in the 5 minutes prior to the OMR assay. A total of 120 larvae per fish line were recorded.

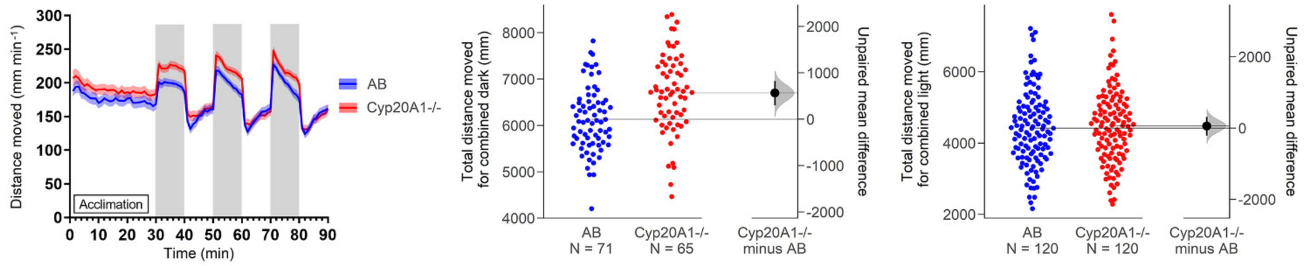


**Figure S3. Locomotion during the light-dark assay of CYP201A1 (wh[61]) mutant larvae in comparison to AB.** CYP201A1 (wh[61]) mutant larvae show hyperactivity in the dark phase but not in the light phase. The experiment was repeated five times independently (total *n* = 120).

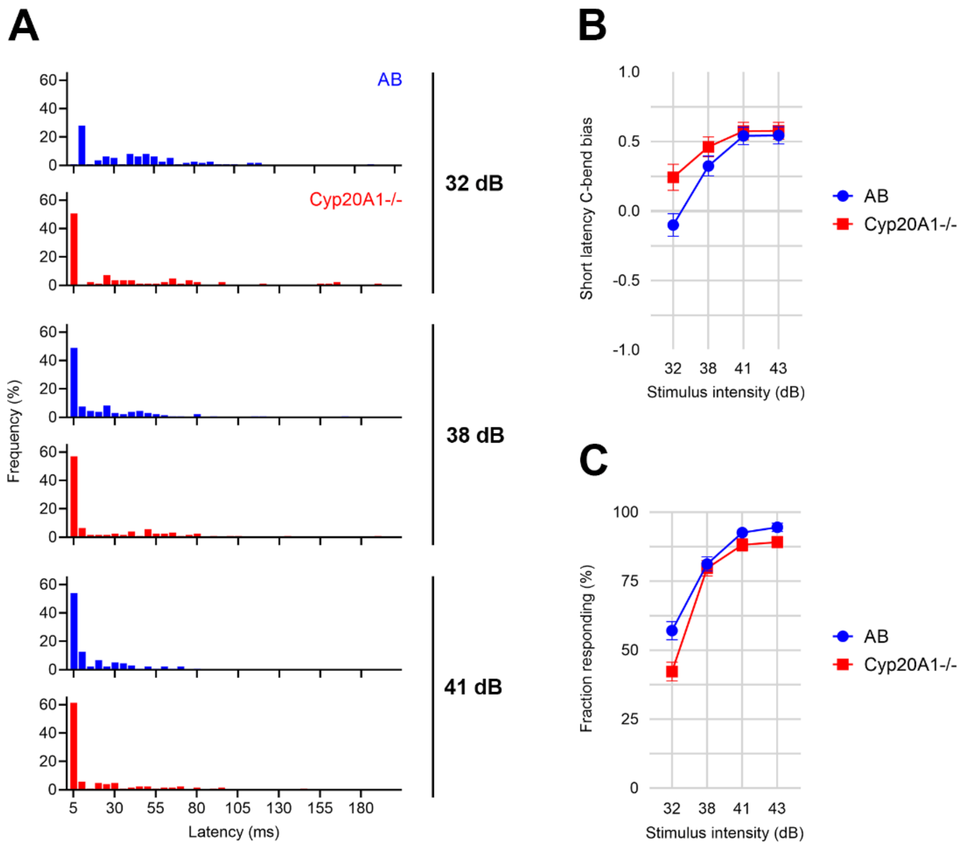


**Figure S4. Startle response.** (**A**) Latency of AB and Cyp20A1-/- wh[61] mutant larvae at 32 dB, 38 dB, and 41 dB. For 43 dB see Figure 2. (**B**) Bias toward short-latency C-bend (< 15 ms) at all stimulus intensities. (**C**) Fraction of larvae responding at different stimulus intensities. All data from three independent experiments.

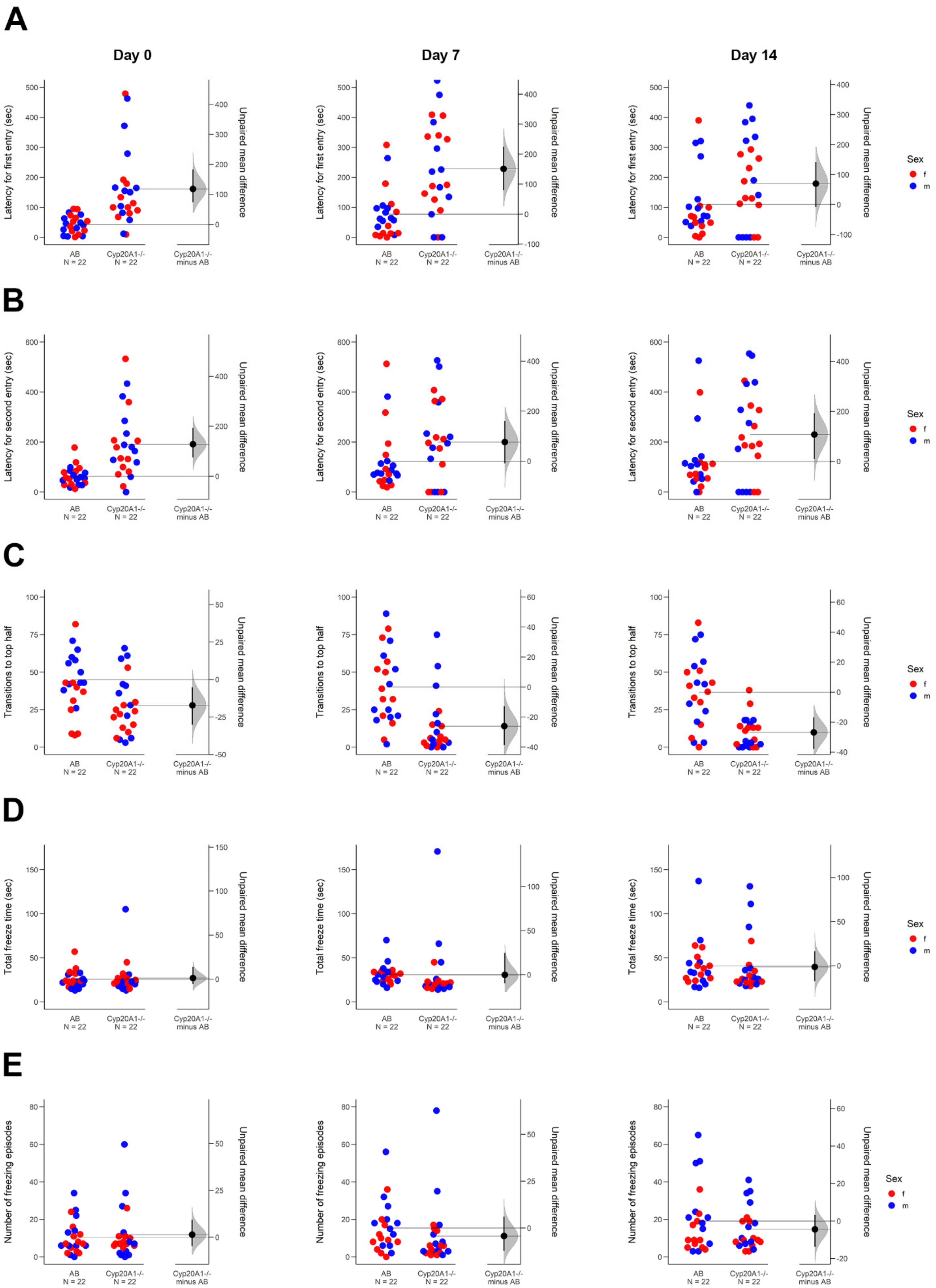
A
B
C
D
E
Day 0
Day 7
Day 14
Latency for first entry (sec)
Latency for second entry (sec)
Transitions to top half
Total freeze time (sec)
Number of freezing episodes
Unpaired mean difference
AB
N = 22
Cyp20A1-/-
N = 22
Cyp20A1-/-
minus AB
Sex
f
m

**F**

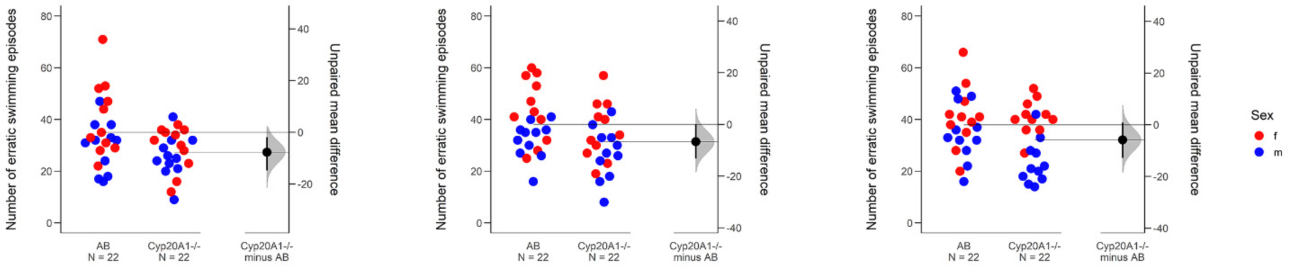

**Figure S5. Adult behavior in the novel tank assay.** (**A**) Latency for first entry to the top half, (**B**) latency for second entry to the top half, (**C**) number of transitions to the top half, (**D**) total freeze time, (**E**) number of freezing episodes, (**F**) number of erratic swimming episodes (also called darting). The experiment was repeated on day 7 and day 14 with the same fish.
